# Locus-Level Transposable Element Profiling Resolves Division-Coupled Transcriptional Dynamics During Human Endoderm Specification

**DOI:** 10.64898/2026.07.07.737093

**Authors:** Amrit Gaire, Erich Kummerfeld, Constantin Aliferis, Jinhua Wang

## Abstract

**Background:** Transposable elements (TEs) constitute nearly half of the human genome and are now recognized as significant contributors to mammalian gene regulatory networks. Despite this, most transcriptomic studies quantify TE expression at the subfamily level, which may obscure meaningful variation arising from individual insertion sites. Whether resolving TE expression to individual loci can reveal biologically distinct signals during stem cell differentiation has not been systematically characterized.

**Results:** We re-analysed a published RNA-seq time course of FUCCI-h9 human embryonic stem cells differentiating into definitive endoderm (0–72 hour, seven time points, three division cycles, three biological replicates), quantifying expression in parallel at two complementary resolutions: TE subfamilies using TEtranscripts and individual TE loci using TElocal. The primary finding is that individual TE loci capture heterogeneous transcriptional responses at cell division boundaries that are entirely absent at the subfamily level. Within-division-state PC1 variance for TE loci was substantially elevated at the first division cycle (6.72; 95% bootstrap CI 0.63–8.48) compared with TE subfamilies (0.22; CI 0.04–0.34), with non-overlapping confidence intervals providing statistically robust support for the resolution advantage. Differential expression analysis identified over 18,000 dynamic TE loci across the time course, exceeding the 268 differentially expressed subfamilies, with alternating phases of silencing and reactivation resolved only at locus resolution. Differentially expressed TE loci were non-randomly enriched at superenhancers (fold enrichment: 9.1–23.5-fold; p < 0.001 by permutation), with peak overlap at 36–48 hours coinciding with the second cell division and endoderm commitment. ERV1-class elements, particularly HERVH-int, were the dominant contributors, and representative loci near the endoderm regulators MIXL1 and ID3 showed differentiation-induced RNA-seq signal within proximal superenhancer domains.

**Conclusions:** TE loci exhibit heterogeneous transcriptional responses at cell division boundaries, a signal with non-overlapping bootstrap confidence intervals relative to TE subfamilies at the first division cycle that is entirely masked by subfamily-level aggregation. This division-boundary resolution advantage, together with permutation-confirmed enrichment of dynamic loci at superenhancers during a discrete 36–48 hour endoderm commitment window, supports broader adoption of locus-resolved TE quantification as a complement to conventional gene expression analysis in developmental genomics.

## Introduction

Transposable elements (TEs) are mobile genetic elements that make up close to half of the human genome. They were historically dismissed as non-functional repetitive sequence, though it is now clear that TE-derived sequences make substantial contributions to gene regulation across mammalian cell types^1–4^. TE sequences can function as promoters, enhancers, silencers, and insulators^4^, and their expression is frequently tissue-specific^5^. The RNA products of TEs have been linked to epigenetic programs underlying cell specification and differentiation^6,7^. In pluripotent stem cells and during early embryogenesis, evolutionarily young TE families act as cis-regulatory elements that shape transcriptional networks and influence chromatin architecture^4^. HERVH and LTR7 elements, for example, have been reported to function as OCT4 and NANOG-bound enhancers and alternative promoters for a subset of pluripotency-associated genes^4,8^.

Early lineage specification is a fundamental phase of embryonic development, producing the three primary germ layers from which all tissues eventually arise^9^. During directed differentiation of human pluripotent stem cell (hPSCs) toward these lineages, TE show lineage-specific expression patterns and dynamic subcellular localization^10^. TE-containing transcripts showed cell type specific stability profiles and increased chromatin association in differentiated lineages^10^, pointing to possible roles in post-transcriptional regulation during germ layer specification. Beyond this general involvement, specific TE subfamilies such as LTR7 elements have been shown to act as mechano-responsive enhancer elements that inhibit definitive endoderm (DE) differentiation in hESCs through regulation of genes including FAM189A2^11^. A recent study also showed that key differentiation genes are transcribed and enhancers are established before cell division occurs, raising the possibility that TE-associated regulatory elements participate in priming gene expression programs prior to mitosis and contribute to endoderm specification^12^.

One persistent limitation in the field is that most studied aggregate TE expression at the family or subfamily level. This aggregation may mask locus-specific heterogeneity arising from insertion site-specific regulation^13,14^. Tools capable of quantifying TE expression at the individual locus resolution have begun to address this, but systematic comparisons of subfamily-level versus locus-level analysis during continuous developmental trajectories, and their relationship to cell division-driven transcriptional transitions, remain largely unexplored. To begin filing this gap, we reanalysed publicly available RNA-seq data from a time course study spanning 0 to 72 hours of endoderm differentiation^12^. The dataset includes seven time points with three biological replicates per condition, capturing transcriptional changes across both chronological time and three successive cell divisions. Figure 1a summarizes the experimental design of the original study, including sampling time points and division stages. We used a computational workflow that quantifies TE expression at both the subfamily level using TEtranscripts^15^ and at individual locus resolution using TElocal^15,16^ (Figure 1b), with the same underlying alignment framework for both. This parallel design allows direct comparison of the two resolution levels from the same RNA-seq data.

**Figure 1.**
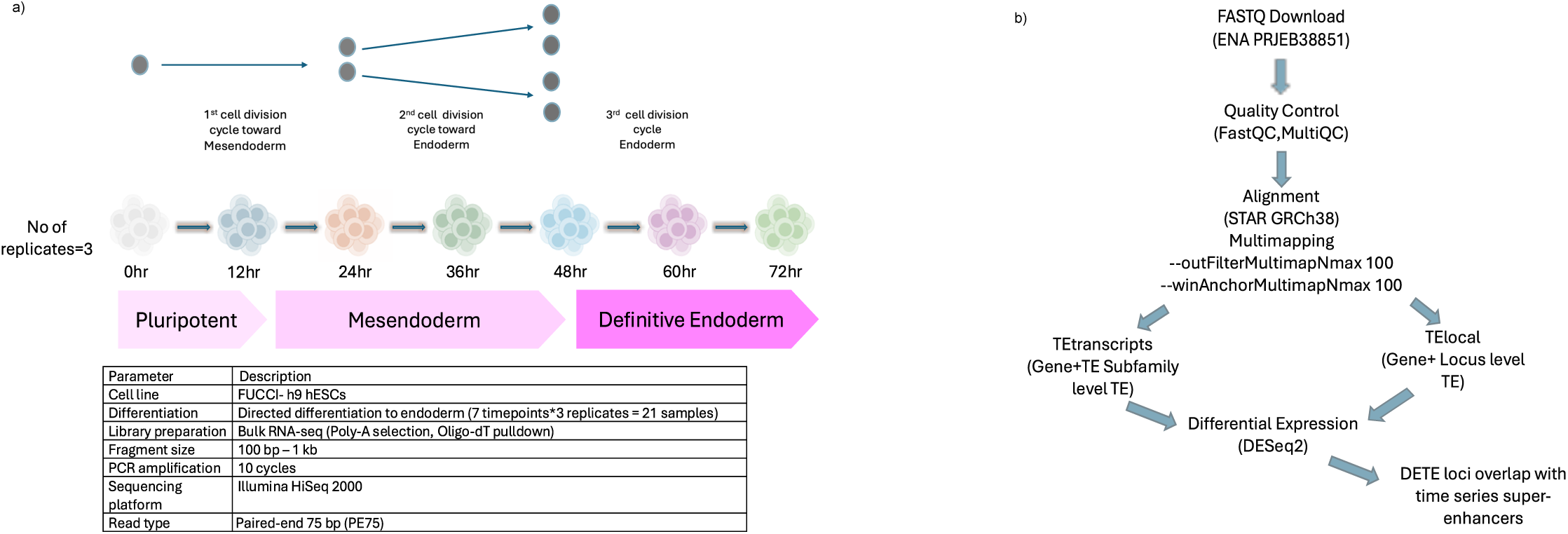
Experimental design and computational workflow. (a) FUCCI-h9 human pluripotent stem cells were directed toward definitive endoderm and sampled at seven time points (0, 12, 24, 36, 48, 60, and 72 hours) spanning three successive cell-division cycles, with three biological replicates per time point (21 samples total; ENA accession PRJEB38851). The table summarizes key experimental parameters. (b) RNA-seq and TE analysis pipeline: raw FASTQ files underwent quality control, alignment to GRCh38 with STAR (multi-mapping allowed), gene and TE subfamily quantification with TEtranscripts, locus-specific TE quantification with TElocal, differential expression with DESeq2, and intersection of DETE loci with time-series superenhancers visualised in IGV.

Our analysis shows that individual TE loci track differentiation progression more tightly than either coding genes or TE subfamilies do, and that locus-level data captures heterogeneous transcriptional responses at division state boundaries that are entirely masked at the subfamily level. In addition, dynamically expressed TE loci are significantly enriched within superenhancers during a discrete 36-to-48-hour window corresponding to second cell division and the onset of endoderm commitment, with specific loci identified near the endoderm master regulators MIXL1 and ID3. We discuss the biological implications of these findings alongside the considerable limitations of a computational reanalysis and propose that locus resolved TE profiling needs a broader adoption as a complement to conventional gene expression analysis in developmental studies.

## Methods

### Data acquisition and quality control

Paired-end RNA sequencing data were obtained from the NCBI Sequence Read Archive (SRA) under accession PRJEB38851 (run accessions ERR4235451–ERR4235471), corresponding to a time-course experiment in human embryonic stem cells (hESCs) sampled at 0, 12, 24, 36, 48, 60, and 72 hours post-differentiation induction, with three biological replicates per time point (21 samples total)^12^. FASTQ files were downloaded using SRA Toolkit v3.0.0^30^ with --split-files option. Read quality was assessed using FastQC v0.11.9^31^, and aggregated MultiQC v1.11^32^.

### Read alignment

Reads were aligned to the GRCh38 primary assembly using STAR v2.7.11b^33^ with a genome index built from Homo_sapiens.GRCh38.113.gtf (--sjdbOverhang 74). Alignment was performed with --outSAMtype BAM SortedByCoordinate and multi-mapping parameters (--outFilterMultimapNma× 100, --winAnchorMultimapNma× 100) to retain reads mapping to repetitive TE sequences^15^. BAM files were indexed using SAMtools v1.13^34^.

### Transposable element quantification

TE expression was quantified at two complementary levels of resolution using tools from the Hammell laboratory ^15,16,35^. Subfamily-level quantification used TEcount from the TEtranscripts suite (--mode multi, --sortByPos) with GRCh38 Ensembl gene and RepeatMasker TE annotations. Locus-level quantification used TElocal (--mode multi, --sortByPos) with a custom TE locus index (GRCh38_Ensembl_rmsk_TE.gtf.locInd).

### Differential expression and statistical modeling

Raw count matrices were imported into R v4.4.0^36^ and low-count features were removed using filterByExpr() from edgeR (v4.4.2)^37^ with sample grouping specified by the primary variable of interest. Filtered matrices were loaded into DESeq2 (v1.46.0)^38^ with design formulae ∼ replicate + division_state or ∼ replicate + time. VST normalisation was applied using vst(dds, blind = FALSE) for primary analyses and vst(dds, blind = TRUE) for sensitivity assessment. Differential expression was called at padj ≤ 0.05 and |log₂FC| > log₂(1.5). Pairwise contrasts were computed relative to the undifferentiated/0 h baseline, between consecutive time points, and across division-state transitions.

### Principal component analysis and clustering metrices

PCA was performed on VST-normalized values using the top 500 most variable features (plotPCA(), DESeq2), with robustness confirmed using 2,000 (genes, TE loci) or 1,000 (TE subfamilies) features. The linear association between PC1 scores and numeric division-state encoding (1–4) or time (0–72 h) was quantified by Pearson correlation and the coefficient of determination (R²) from linear regression. Bootstrap confidence intervals (95%, n = 1,000 resamples) were computed by resampling samples with replacement and re-fitting the linear model. Mean silhouette scores were computed in PC1–PC2 Euclidean space using the silhouette() function from the cluster package (v2.1.8)^39^. Within-state and within-timepoint PC1 variance was calculated across replicate PC1 scores within each grouping level.

### K-means clustering

K-means clustering (k = 4) was applied to the top 100 most variable TE features per resolution level, using z-score-normalized VST expression. The choice of k = 4 was guided by the four biological division states in the experimental design (undifferentiated, 1st cycle, 2nd cycle, 3rd cycle). Cluster membership was held constant between time-ordered and division-state-ordered sample arrangements to allow direct visual comparison.

### Super-enhancer overlap analysis

Genomic coordinates of DETE loci were extracted from the TE locus index (GRCh38_Ensembl_rmsk_TE.gtf.locInd.locations)^16^ and restricted to canonical chromosomes. Superenhancer annotations at each time point (0, 12, 24, 36, 48, and 72 h) were taken from original study^12^ and classified as proximal or distal relative to the nearest TSS.60 hr was excluded because of no superenhancers annotations in the original study. Overlap with DETE loci was computed using findOverlaps() from GenomicRanges package (v1.58.0)^40^ with a minimum overlap of 50 bp. Temporal persistence of overlapping loci was visualised using UpSet plots UpSetR package (v1.4.0)^41^. Genome-browser snapshots were generated in IGV.

### Visualization and software

All plots were generated using ggplot2 (v4.0.2)^42^ with composite figures assembled using patchwork package (v1.3.2)^43^ and cowplot (v1.2.0)^42^. Heatmaps used ComplexHeatmap (v2.22.0)^44^ with colour scales from circlize package (v0.4.16)^45^ and RColorBrewer (v1.1-3)^46^. All analyses were performed under Rocky Linux 8.10 using R v4.4.0 with Bioconductor v3.20^47^.

## Results

### Locus-level TE profiling captures stronger transcriptional structure along the differentiation trajectory

To compare how well protein coding genes, TE subfamilies and individual TE loci reflect differentiation state, we computed variance-stabilized expression (VST; blind = FALSE) across all 21 samples for both division states and timepoints and performed principal component analysis (PCA) on the 500 most variable features from each feature class. Sample-to-sample Pearson correlation heatmaps showed tight within-group clustering and progressive between-group separation for all three feature classes under both division state and chronological time models (Figures 2a–c and 3a–c). Block-diagonal structure sharpened from the undifferentiated state through to the third cell division cycle, and replicate consistency was strong across all conditions.

**Figure 2.**
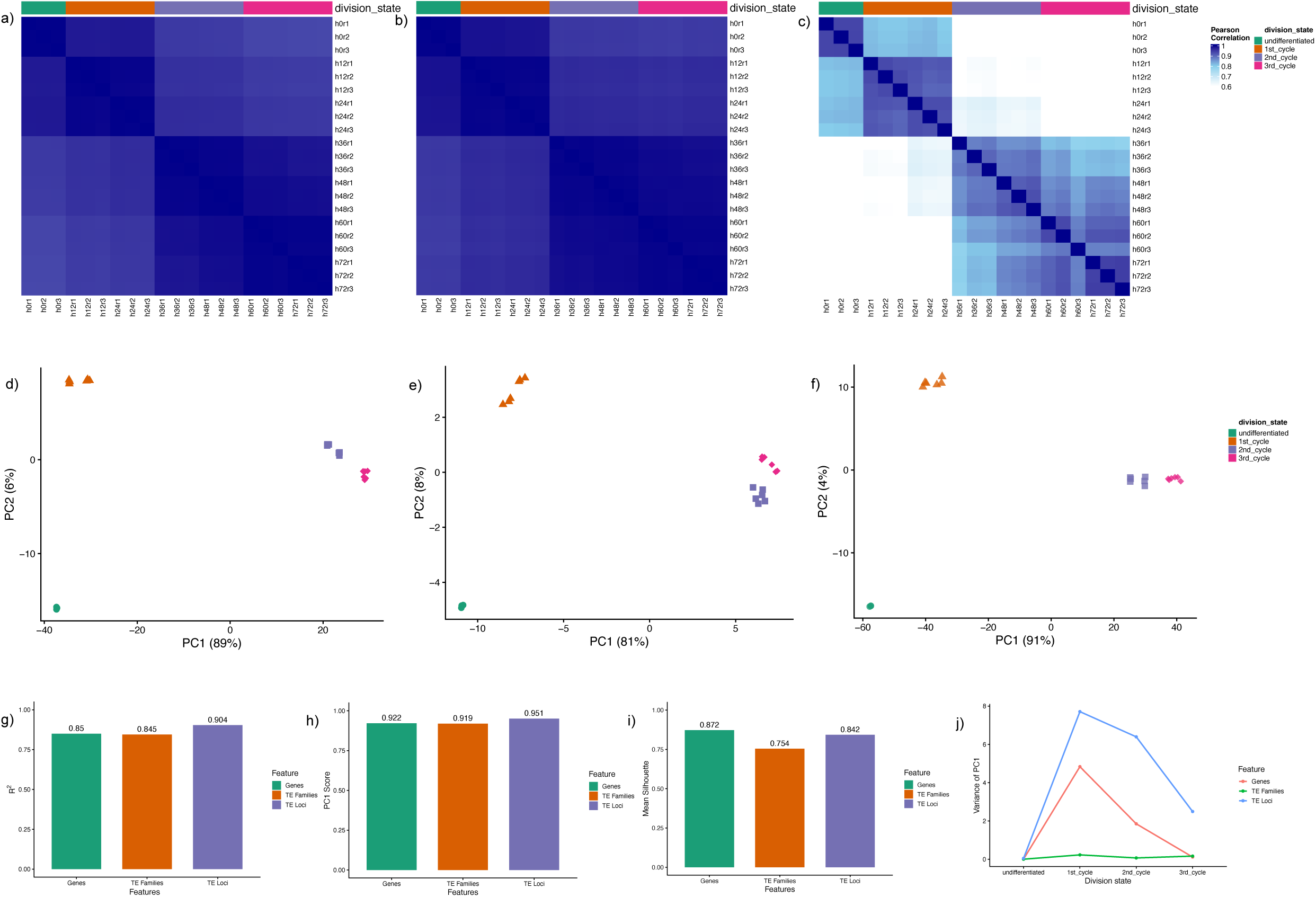
Cell division state structures transcriptional variation in genes and transposable elements. (a-c) Pearson correlation heatmaps of VST-normalised expression for (a) protein-coding genes, (b) TE subfamilies, and (c) individual TE loci, ordered by division state. (d-f) PCA of the top 500 most variable features for (d) genes, (e) TE subfamilies, and (f) TE loci, coloured by division state. (g) R² from linear regression of PC1 scores against numeric division state. (h) Pearson correlation coefficients between PC1 scores and division state. (i) Mean silhouette scores in PC1-PC2 space. (j) Within-division-state variance of PC1 scores; TE loci show markedly elevated variance at the first and second cycle boundaries. Bootstrap confidence intervals are provided in Supplementary Table S3.

**Figure 3.**
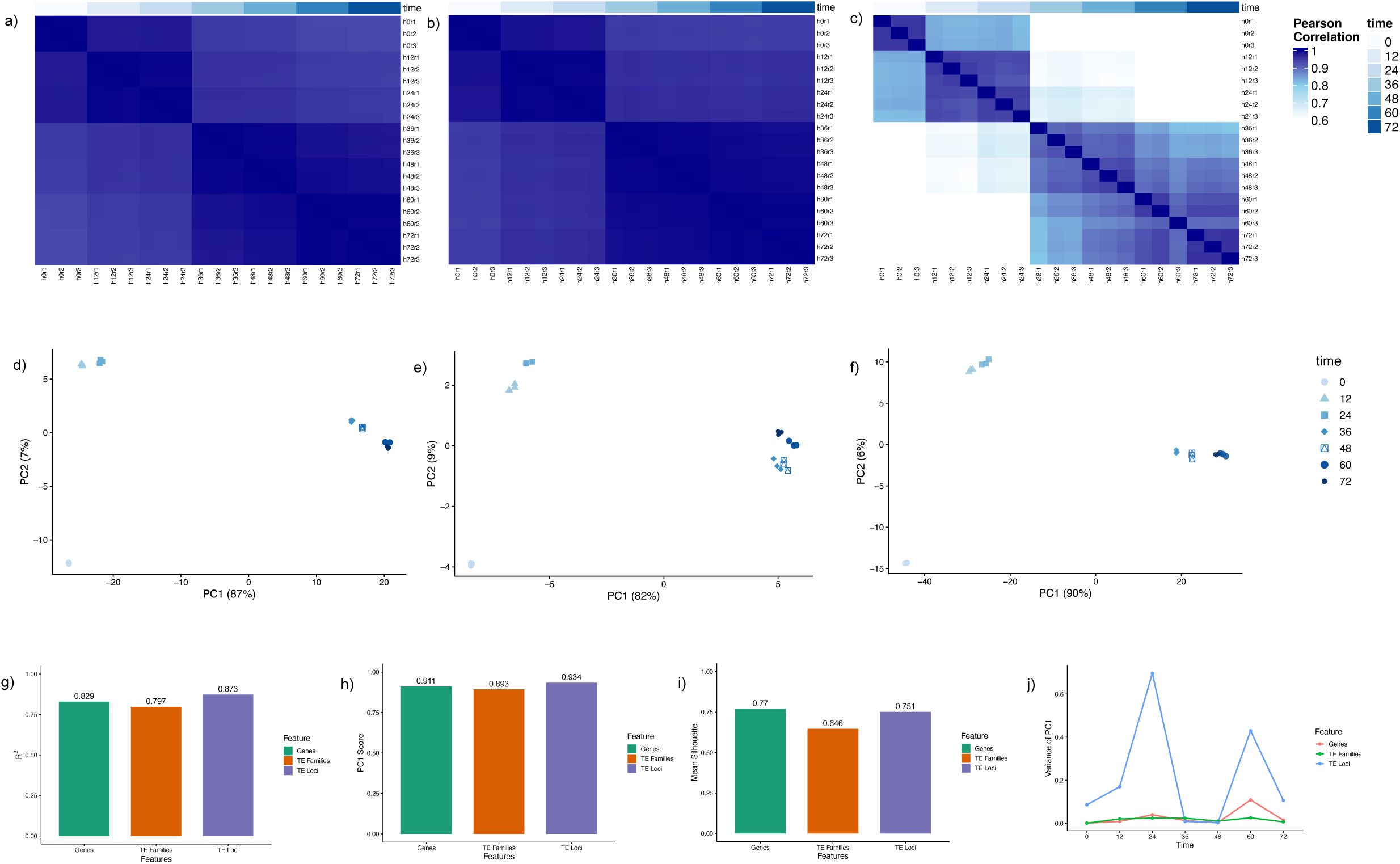
Chronological time captures differentiation progression with lower resolution than division state. (a-c) Pearson correlation heatmaps ordered by chronological time for (a) genes, (b) TE subfamilies, and (c) TE loci. (d-f) PCA of the top 500 most variable features coloured by time point. (g) R² from linear regression of PC1 against chronological time. (h) Pearson correlation of PC1 with time. (i) Mean silhouette scores. (j) Within-time-point variance of PC1 scores; TE loci spike at 24 and 60 hours, corresponding to division-cycle boundaries.

PCA under the division state model (Figure 2d–f) produced a clean linear trajectory along PC1 for all feature classes. PC1 captured 89%, 81%, and 91% of total variance for genes, TE subfamilies, and TE loci respectively, consistent with a single dominant axis of variation in this synchronized system. Linear regression of PC1 scores against numeric division state (undifferentiated = 1 through 3rd cycle = 4) yielded R² values of 0.850, 0.844, and 0.905, with Pearson correlation coefficients of 0.922, 0.919, and 0.951 for genes, TE subfamilies, and TE loci respectively (Figure 2g-h). By both metrics, TE loci tracked differentiation progression most closely among the features examined. Analogous analyses under the chronological time model gave a comparable ordering (R² = 0.829,0.797,0.873 respectively; Figure 3g). However, bootstrap confidence intervals (n=1000 resamples, Supplementary Table S1) showed the advantage of division state over time did not reach statistical significance for any feature class, with all confidence intervals including zero. This is not surprising given the near-collinearity of division state and chronological time in a synchronized FUCCI-hESCs system, and the more informative comparison is within-state variance analysis described below.

Mean silhouette scores in PC1–PC2 space were positive for all three feature classes: 0.872 for genes, 0.754 for TE subfamilies, and 0.842 for TE loci under the division state model (Figure 2i). The lower score for TE subfamilies relative to both genes and TE loci is consistent with the averaging effect of family-level aggregation. R² and Pearson correlation provide a more direct measure of continuous trajectory fit than silhouette score does, and by these metrices TE loci were the most informative feature class. These findings held across different feature selection thresholds (Supplementary Figure 1, using the top 2000 or 1,000 features) and under blind=TRUE VST normalization (Supplementary Figure 2), suggesting the locus-level advantage is not an artefact of the normalization approach. We note that this conclusion resets on a single dataset, and whether same ordering holds across other differentiation systems or cell lines cannot be determined from the present analysis.

### Within-division-state variance of TE loci identifies division-boundary dynamics masked at subfamily resolution

The most distinctive signal in the locus-level data was seen from examining within-state PC1 variance across feature classes (Figure 2j). TE loci showed a pronounced increase in variance during the first division cycle (6.72; 95% bootstrap CI 0.63 to 0.48), which remained elevated in the second cycle (6.02; confidence interval (CI) 0.009 to 6.28). These values were substantially higher than those for TE subfamilies (first cycle: 0.22, CI 0.04 to 0.34; second cycle: 0.07, CI 0.01 to 0.11). Notably, the bootstrap CIs for TE loci and TE subfamilies did not overlap during the first division cycle, providing strong quantitative evidence for the higher resolution at the locus level. By the third cycle, TE loci variance declined (2.01; CI 0.41 to 3.10), coinciding with consolidation of the endoderm transcriptional state. In contrast, TE subfamily variance remained relatively stable across division states, and gene-level PC1 variance was ∼1.4-fold lower than that of TE loci at the first cycle boundary.

Under the chronological time model (Figure 3j), TE loci showed variance spikes at 24 hours (0.697; CI 0.00 to 0.907) and 60 hours (0.430; CI 0.00 to 0.571), time points that correspond to division state boundaries in this synchronized system, whereas genes showed only a modest elevation at 60 hours and TE subfamilies remained low throughout. The wide confidence intervals in the time model reflect the limited statistical power given by three biological replicates per time point, and the point estimates should be interpreted with caution. Nevertheless, the pattern of elevated variance at division boundaries is consistent across both organizational frameworks. Collectively, these within-state variance data suggest that individual TE loci undergo heterogeneous transcriptional responses at cell cycle boundaries, a dynamic signal that appears to be averaged away when expression is aggregated across all insertions of a given subfamily. Because this analysis is conducted within each grouping level independently, it is not confounded by the collinearity of division state and time noted above.

### Global TE expression dynamics reveal structured waves of silencing and reactivation across differentiation

To characterize global TE expression dynamics, we computed mean VST expression across all TE subfamilies and loci at each time point and division state (Figure 4a). Mean subfamily level expression declined sharply between 0 and 12 hr, recovered gradually, peaked at 48hr and plateaued thereafter (Figure 4a-i). At individual locus resolution, the decline was more gradual, reaching a minimum at 36 hr before partial recovery toward 72 hr (Figure 4 a-iii). When sample were ordered by division state, subfamily level expression was lowest at the first division cycle and partially recovered thereafter, whereas locus level expression reached its minimum at the second division cycle (Figure 4a-ii, iv). The later and more prolonged silencing detected at locus resolution compared with subfamily resolution is consistent with the locus level approach capturing insertion specific dynamics that are averaged out at the family level.

**Figure 4.**
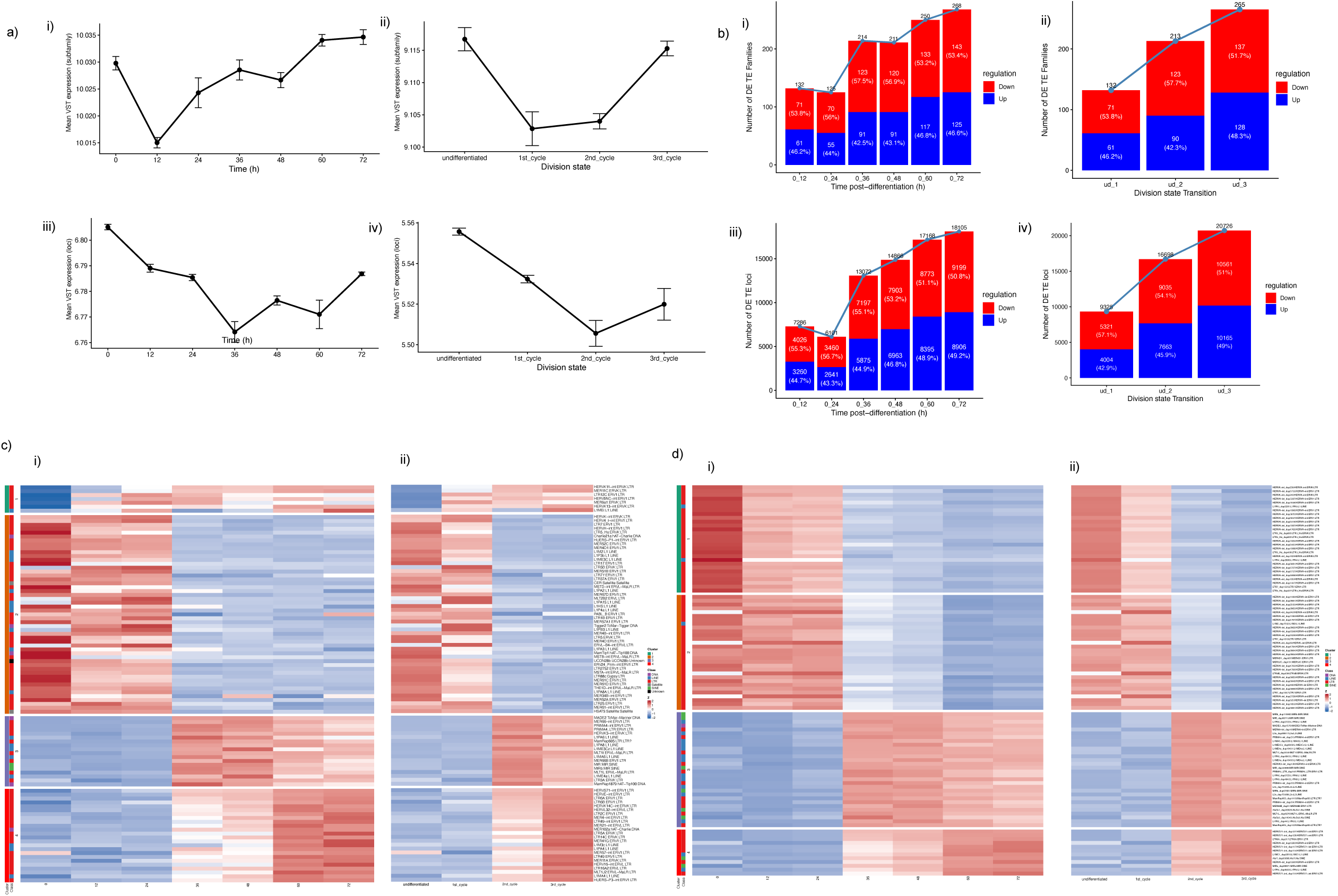
TE expression dynamics reveal structured waves of silencing and reactivation. (a) Mean VST expression of TE subfamilies (i, ii) and individual TE loci (iii, iv) across chronological time (i, iii) or division state (ii, iv). Points represent means of three biological replicates; error bars show SEM. (b) Number of significantly differentially expressed TEs (padj ≤ 0.05, |log₂FC| > log₂(1.5)) at each time point (i, iii) or division-state transition (ii, iv) for TE subfamilies and loci. (c) Z-score heatmaps of the 100 most variable TE subfamilies ordered by time (i) or division state (ii); rows grouped by k-means clustering (k = 4). (d) As (c) but for individual TE loci; HERVH-int loci dominate cluster 1, and Alu-specific loci are resolved only at locus resolution.

Differential expression analysis (DESeq2; padj ≤ 0.05 and |log_2_FC| > log_2_(1.5), relative to undifferentiated/0h baseline) showed a progressive accumulation of differentially expressed TE elements over the time course, with downregulation predominating at all time points (Figure 4b). At the subfamily level, the number of differentially expressed (DE) subfamilies declined slightly between 12 and 24 hr (132 to 125) before rising sharply from 36 hr onward, reaching 268 at 72 hr (Figure 5b-i). At locus resolution, the same transient reduction was observed at 24 hr (7286 to 6101 DE loci), followed by sharp increase to 13072 at 36 hr and continued accumulation to 18105 at 72 hr (Figure 4b-iii). The proportion of downregulated elements decreased progressively with time, from approximately 57% at 36 hr to 51% at 72 hr for loci, reflecting a gradual increase in the upregulated fraction as differentiation proceeds rather than a directional reversal. Comparable trends were observed across division state transitions (Figure 4b-ii, iv).

**Figure 5.**
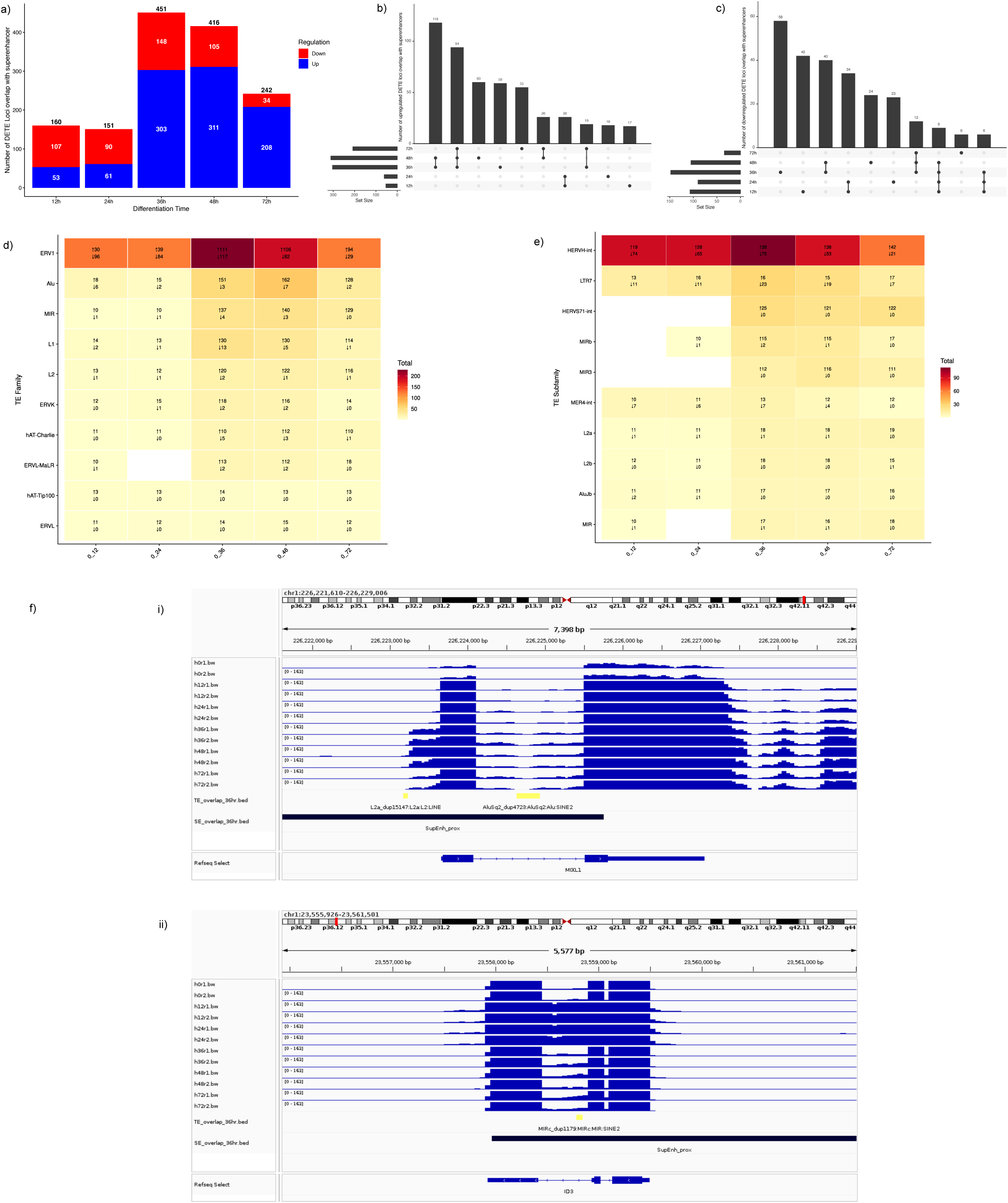
Dynamic TE loci are non-randomly enriched at superenhancers near endoderm master regulators. (a) Number of DETE loci overlapping superenhancers (minimum 50 bp) at each time point, split by upregulation and downregulation. (b, c) UpSet plots of superenhancer-overlapping upregulated (b) and downregulated (c) DETE loci across the time course. (d, e) Heatmaps of the top TE families (d) and subfamilies (e) contributing to superenhancer-overlapping DE loci. (f) Genome browser snapshots of (i) L2a LINE (L2a_dup15147) and AluSq2 SINE (AluSq2_dup4723) within the MIXL1 proximal superenhancer (chr1:226,221,610-226,229,086), with signal peaking at 36 to 72 hours; and (ii) MIRc SINE (MIRc_dup1179) within the ID3 proximal superenhancer (chr1:23,555,926-23,561,501), with progressive signal accumulation from 12 hours. Minimal signal at 0 hours confirms differentiation-induced activation in both cases.

K means clustering (k=4, selected to correspond to the four biological division states as a confirmatory rather than exploratory framework) of the top 100 most variable TE features highlighted a key distinction between subfamily and locus resolution (Figure 4 c-d). At the subfamily level, all four clusters were compositionally heterogeneous, with no single TE class dominating any cluster (Figure 4c-i). Cluster 1 and 2 contained primarily ERVK and ERV1 LTR elements including HERVK int, LTR5_Hs, LTR7 and HERVH-int, whereas Cluster 3 and 4 contained a mixture of older ERVL-MaLR, ERVL and LINE elements showing more gradual early repression. At locus resolution, Cluster 1 was dominated by HERVH-int loci, with smaller contributions from HERVK-int, LTR7 and LTR5_Hs loci, consistent with the known abundance of HERVH transcripts in human embryonic stem cells and their established role as markers of pluripotency^17^. Notably, HERVH-int loci also contributed disproportionately to the early repressive wave, a pattern masked when expression is averaged at the subfamily level. Specific Alu loci (AluSx1, AluY and AluSx3) appeared only in the locus level clusters, indicating insertion site specific regulation that is not reflected in subfamily level summaries. Reordering samples by division state sharpened cluster boundaries at both resolutions, converting gradual temporal gradients into discrete, stable blocks (Figure 4c ii, 4d ii).

Stepwise analysis between consecutive timepoints revealed alternating phases of high and low TE transcriptional activity (Supplementary figure 3 a-c). The 0-12 hr and 24-36 hr intervals were the most dynamic, yielding 7286 and 9253 DE loci and 132 and 154 DE subfamilies respectively with downregulation predominating (∼55-60%) at both. Intervening near silent windows at 12-24 hr (329 DE loci, 6 subfamilies) and 36-48 hr (597 loci, 7 subfamilies) were enriched for upregulation (71% and 86% respectively), representing a pattern opposite to that seen during the active phases. Across division state transitions, DE counts peaked at the 1^st^ to 2^nd^ cycle step (15196 loci, 169 subfamilies), with downregulation dominant (Supplementary Figure 3 d-f). At the 2^nd^ to 3^rd^ cycle transition, total DE counts fell sharply (4719 loci, 55 subfamilies) and directionality reversed, with upregulation becoming the majority for loci (57.9%) and subfamilies (72.7%), suggesting partial reactivation of TE elements following the completion of the main repressive phase. Genome wide heatmaps of highly significant DE elements showed that division state ordering produced cleaner cluster boundaries at both resolutions, converting gradual temporal gradients into discrete, stable expression blocks (Supplementary Figure 3g i-iv).

### Dynamic TE loci are non-randomly enriched at superenhancers during endoderm commitment

To assess the potential regulatory significance of dynamically expressed TE loci, we intersected DETE loci (DETE loci; minimum overlap of 50 bp) with superenhancers mapped across the pluripotency to definitive endoderm time course from original study^12^. The 60 hour timepoint was excluded because no superenhancers annotations were reported at that time point. Permutation testing (1000 random coordinate shuffles restricted to canonical chromosomes) confirmed that the observed overlaps are significantly greater than expected by chance at all analysed time points (Table 1). Fold enrichment over the permutation null ranged from 9.1-fold at 36 hours to 23.5-fold at 24 hours, with all p-values below 0.001.

**Table 1.** Enrichment of differentially expressed TE loci at superenhancers (permutation test, n = 1,000 shuffles)

| Timepoint | Observed overlaps | Null mean | Null 95% CI | Fold enrichment | p-value | Sig. |
| --- | --- | --- | --- | --- | --- | --- |
| 12 h | 160 | 8.44 | 3-15 | 18.97x | <0.001 | *** |
| 24 h | 151 | 6.44 | 2-11 | 23.45x | <0.001 | *** |
| 36 h | 451 | 49.34 | 37-63 | 9.14x | <0.001 | *** |
| 48 h | 416 | 33.89 | 23-46 | 12.27x | <0.001 | *** |
| 72 h | 242 | 20.45 | 12-30 | 11.83x | <0.001 | *** |

Note: Permutation null was derived from 1,000 random coordinate shuffles of DETE loci across canonical chromosomes. The 60-hour time point was excluded due to absence of superenhancer annotations in the source dataset.

The number of overlapping DETE loci increased progressively, peaking at 36-48 hr with 451 and 416 overlapping loci respectively, before declining to 242 loci at 72hr (Figure 5a). Upregulated loci consistently outnumbered downregulated after early time points, particularly at 36-48 hr. Upset analysis showed the majority of superenhancers overlapping loci were time point specific with 36 hr and 48 hr sharing the largest intersections among both upregulated (118 loci) and downregulated (40 loci) sets (Figure 5 b-c), consistent with a discrete window of TE-superenhancer coincidence during the second cell division cycle.

At the family level ERV1 class elements were the largest contributors to superenhancers overlapping loci at all time points, with peak activity at 36-48 hr (Figure 5d). At subfamily resolution, HERVH-int was the single largest contributor throughout the time course, followed by LTR7, consistent with the established roles of these elements as regulatory hubs in pluripotent and early lineage committed cells^8^ (Figure 5e). Notably, MIR and Alu loci which were near absent from the superenhancers overlap at early timepoints were seen accumulating progressively from 36hr onward and were mostly upregulated. This shows a distinct cluster of evolutionarily older TE loci that may contribute to endoderm specific enhancer activity independently of the HERVH/LTR7 axis.

Genome browser inspection of representative loci shows the spatial relationship between DETE loci and superenhancers boundaries (Figure 5f). At the MIXL1 locus, an L2a LINE element (L2a_dup15147) and an AluSq2 SINE (AluSq2_dup4723) overlapped the proximal superenhancer domain, with RNA-seq signal peaking at 36 hr across biological replicates and sustained through 72 hr. MIXL1 encodes a paired-type homeodomain transcription factor expressed at the primitive streak that is required for mesendoderm patterning^18,19^, making this loci a canonical marker of the endoderm commitment transition. At the ID3 locus, a single MIRc SINE (MIRc_dup1179) was identified within a broad proximal superenhancer domain and displayed progressive signal accumulation from 12 hr onward, with maximal coverage at 36-48 hr. Study suggests that temporal activation of BMP–SMAD1/5 signaling, which peaks at 36 hr of differentiation and directly induces ID1 and ID3 expression, is required for commitment to the anterior primitive streak and mesendodermal fate in human iPSCs^20,21^. In both cases, RNA-seq coverage at the TE-containing superenhancer regions was minimal at 0 hr, indicating that the observed signal is differentiation induced rather than constitutive. The spatial coincidence of DETE loci with superenhancers boundaries at these two loci, reproduced across independent biological replicates, is consistent with the possibility that these insertions mark transcriptionally active regulatory elements. Whether the TE insertions contribute directly to superenhancers function or whether their upregulation reflects transcriptional read through from the surrounding regulatory architecture cannot be determined from these data alone and require direct experimental investigation.

## Discussion

The central finding of this study is that individual TE loci provide a more granular readout of transcriptional dynamics during human PSC differentiation to definitive endoderm than subfamily-level aggregation does. Three complementary lines of evidence support this conclusion. First, TE loci achieve the highest R² (0.905) and Pearson correlation (r = 0.951) with differentiation progression among the three feature classes examined, reflecting tighter linear correspondence between individual insertion activity and developmental state. Second, within-division-state PC1 variance for TE loci is substantially elevated at the first and second division boundaries, with non-overlapping bootstrap confidence intervals relative to TE subfamilies at the first division cycle. This indicates that individual TE insertions respond heterogeneously at pluripotency exit, a response that is completely averaged away at the subfamily level. Third, locus-level analysis identifies more than 18,000 significantly dynamic TE loci alongside a structured pattern of silencing and reactivation, and a discrete 36 to 48 hour window of TE-superenhancers co-localization confirmed by non-random by permutation testing.

The within-state variance is the clearest finding of the resolution advantage, because it is not affected by the near-collinearity of division state and chronological time that limits interpretation of the overall R² comparisons. The bootstrap confidence intervals for the R² differences are wide and all include zero, meaning that the advantage of division state over time as an organizing variable does not reach statistical significance in any feature class, an honest limitation that we acknowledge. The within-state and within-time-point variance analyses sidestep this problem by operating within each grouping level independently, and the elevation of TE loci variance at division boundaries is robust across both organizational frameworks. This is consistent with the growing evidence that cell division functions as a molecular organizer that imprints transcriptional identity through mitotic bookmarking and replication-coupled chromatin remodelling^12,22–24^. Individual TE insertions, because of their specific genomic contexts and proximity to cis-regulatory elements, may be particularly sensitive reporters of the chromatin remodeling events occurring at pluripotency exit. This is speculative; however, we acknowledge that the data presented here do not allow us to distinguish between TEs as active participants versus passive readouts of these processes.

The enrichment of DETE loci within superenhancers at 36 to 48 hours is biologically coherent with the timing of endoderm specification. HERVH-int elements dominate superenhancer-overlapping DETE loci throughout the time course, consistent with prior work showing that HERVH elements overlap annotated superenhancers in human ESCs, demarcate topologically associating domain boundaries, and interact with Mediator components and CTCF in pluripotent cells^8,25^. The progressive accumulation of MIR and Alu loci at superenhancers from 36 hours onward extends this picture by revealing what appears to be a temporally distinct, evolutionarily older layer of TE regulatory activity emerging at endoderm commitment. MIR elements are among the most ancient TEs in the mammalian genome and have been broadly co-opted as regulatory sequences across tissue types^4^; their appearance at endoderm superenhancers during the second division cycle is an interesting observation that warrants further investigation, though we stress it remains purely correlative at this stage.

The locus-specific examples at MIXL1 and ID3 are useful illustrations of the genome-wide findings. MIXL1 is a well-established marker of mesendoderm whose activation at 36 hours in this dataset corresponds to the known timing of primitive streak specification^12,18,19^. The L2a and AluSq2 loci within the MIXL1 proximal superenhancer show matched temporal induction. Similarly, ID3 activation by BMP-SMAD1/5 signaling marks commitment to anterior primitive streak fate^20,21^,and the MIRc SINE within its proximal superenhancer accumulates signal progressively from 12 hours onward. These examples are suggestive, but we are aware that identifying two loci near known endoderm genes is not sufficient to establish a general pattern, and that spatial co-occurrence within a superenhancers boundary does not demonstrate regulatory function. Whether the identified TE insertions contribute directly to superenhancers function – through transcription factor recruitment, chromatin looping, or non-coding RNA scaffolding – or whether their upregulation is a secondary consequence of superenhancers driven transcriptional read through, remains an open question that requires direct experimental investigation.

Several limitations of the present study should be acknowledged and provide directions for future work. First, all analyses are based on a single publicly available dataset from one differentiation protocol and one hESC line (FUCCI-h9); whether the division state signature generalizes across lineages, cell lines or differentiation conditions remains to be established. Second, the regulatory role of DETE loci within superenhancers is inferred entirely from genomic overlap and RNA seq coverage; no perturbation experiments were performed, and causal contributions cannot be claimed from these data. Finally, while stringent statistical thresholds were applied (padj ≤ 0.05, |log_2_FC|>1.5), single cell resolution or longer time courses could further resolve division cycle dependent dynamics from stochastic transcriptional variability.

Despite these limitations, the study has some practical implications worth noting. Precise monitoring of iPSC differentiation quality remains a bottleneck in regenerative medicine^26^. Because individual TE loci provide a higher resolution readout of division history than coding genes or subfamily averages, they may serve as sensitive indicators of differentiation state in applications where precise staging of iPSC-derived cells matters, such as quality control of clinical grade differentiation protocols. This extension should go through validation across multiple iPSC lines and protocols would be needed before drawing firm conclusions. The same analytical framework may also be applicable in contexts where TE derepression^27^ and superenhancers dysregulation converge^28,29^, such as ageing and cancer, although such extensions would require dedicated studies.

In summary, locus-level TE profiling of a human endoderm differentiation time course reveals transcriptional dynamics that are not visible at the subfamily level: heterogeneous division-boundary responses, alternating waves of silencing and reactivation, and a discrete 36 to 48 hour window of TE-superenhancer co-localization near endoderm master regulators. We interpret these findings as motivation for broader adoption of locus-resolved TE quantification in developmental genomics, while acknowledging that the computational observations reported here represent a starting point rather than a conclusion, and that experimental follow-up is essential.

## Conclusions

Locus-level TE expression analysis during human pluripotent stem cell differentiation to definitive endoderm reveals transcriptional dynamics that are not detectable at the subfamily level, including heterogeneous responses at division cycle boundaries, alternating waves of silencing and reactivation, and enrichment of dynamic loci within superenhancers near endoderm master regulators during a discrete 36 to 48 hour window. Bootstrap confidence intervals confirm that TE loci within-state variance at the first division cycle substantially exceeds that of TE subfamilies, providing quantitative support for the resolution advantage. Permutation testing confirms that DETE loci-superenhancer overlap is non-random (fold enrichment 9 to 24-fold; p < 0.001) and peaks during the second cell division cycle. These findings are based on a single dataset and require replication, and the regulatory significance of the identified loci remains to be tested experimentally. Locus-resolved TE profiling represents a useful complement to conventional gene expression analysis in developmental genomics and warrants broader adoption in studies of lineage commitment and TE-associated regulatory evolution.

## Supporting information

Supplementary Figure 1. PCA robustness using top 2,000 or 1,000 most variable features.

Supplementary Figure 2. PCA and summary metrics under blind = TRUE VST normalization.

Supplementary Figure 3. Stepwise differential expression across consecutive time points and division-state transitions.

## Declarations

## Ethics approval and consent to participate

Not applicable.

## Availability of data and materials

All the codes used for preparation of the manuscript will be available on Github.

## Supplementary Tables

**Supplementary Table S1.**
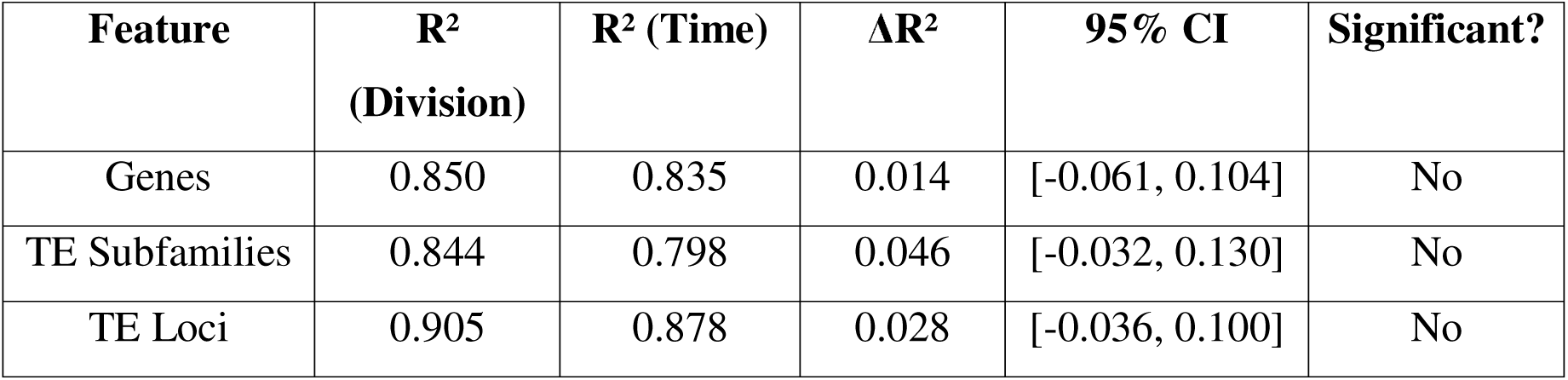
Bootstrap confidence intervals for R² differences (top 500 features)

**Supplementary Table S2.** Within-division-state PC1 variance with bootstrap 95% confidence intervals.

| Feature | Division State | Observed Variance | 95% CI Lower | 95% CI Upper |
| --- | --- | --- | --- | --- |
| Genes | Undifferentiated | 0.006 | 0.000 | 0.006 |
|  | 1st cycle | 4.872 | 0.061 | 5.463 |
|  | 2nd cycle | 1.872 | 0.031 | 2.069 |
|  | 3rd cycle | 0.118 | 0.013 | 0.205 |
| TE Subfamilies | Undifferentiated | 0.003 | 0.000 | 0.004 |
|  | 1st cycle | 0.224 | 0.039 | 0.344 |
|  | 2nd cycle | 0.068 | 0.012 | 0.105 |
|  | 3rd cycle | 0.162 | 0.025 | 0.208 |
| TE Loci | Undifferentiated | 0.061 | 0.000 | 0.066 |
|  | 1st cycle | 6.718 | 0.628 | 8.476 |
|  | 2nd cycle | 6.020 | 0.009 | 6.278 |
|  | 3rd cycle | 2.010 | 0.406 | 3.103 |

Note: TE loci CIs do not overlap with TE subfamily CIs at the first division cycle, supporting the resolution advantage at locus level. n = 1,000 bootstrap resamples.

