## Supplementary figures and images for "Locus-Level Transposable Element Profiling Resolves Division-Coupled Transcriptional Dynamics During Human Endoderm Specification"

### Supplementary Figure 1. PCA robustness using top 2,000 or 1,000 most variable features.

Supplementary Figure 1

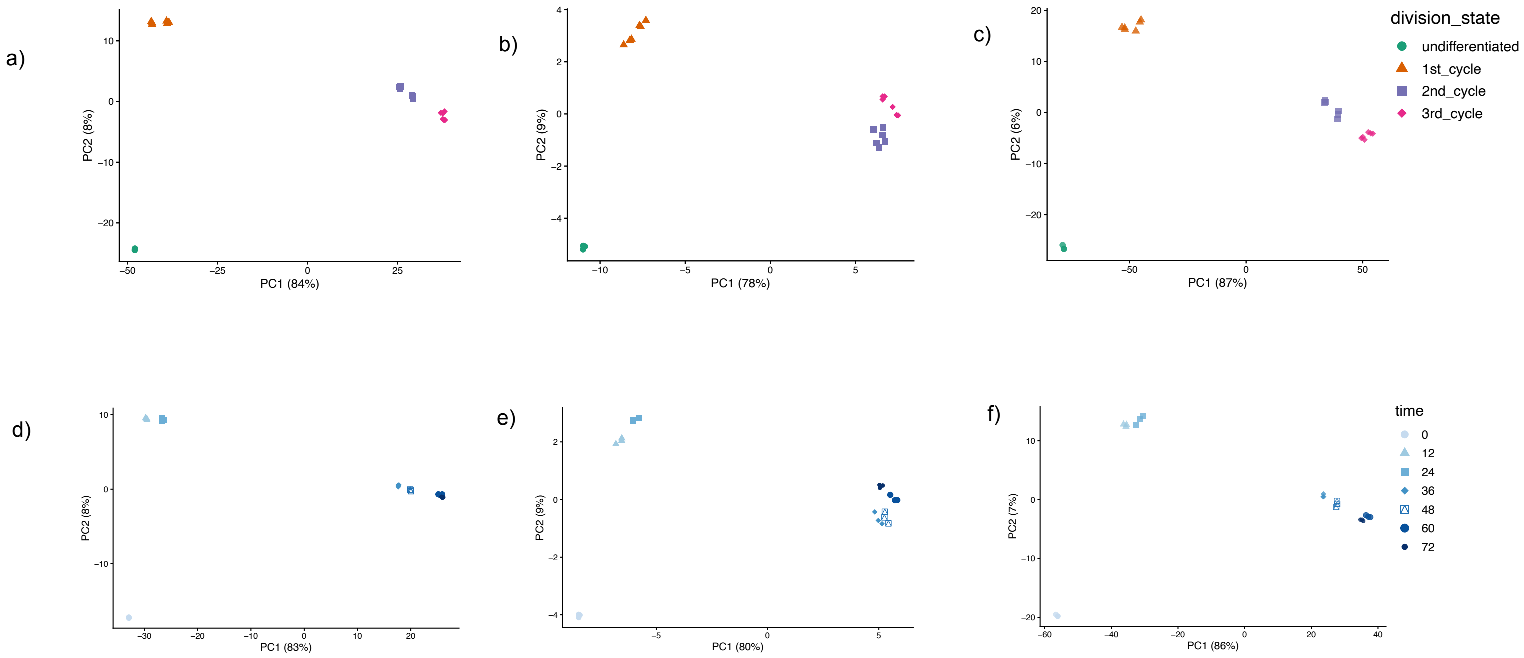

### Supplementary Figure 2. PCA and summary metrics under blind = TRUE VST normalization.

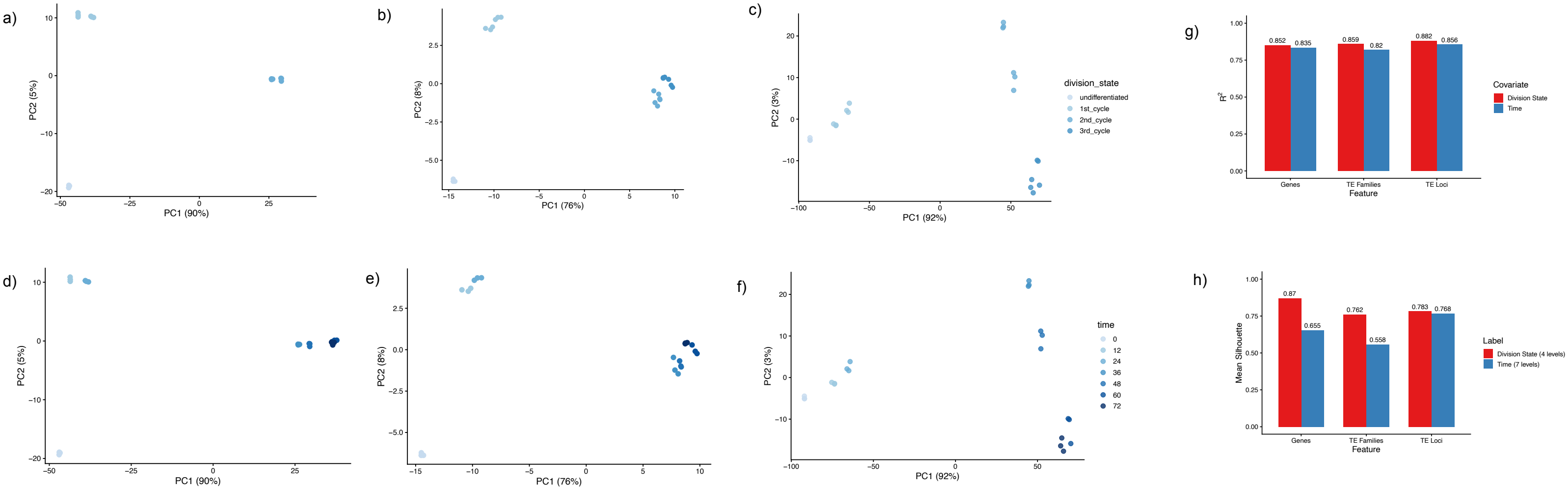

### Supplementary Figure 3. Stepwise differential expression across consecutive time points and division-state transitions.

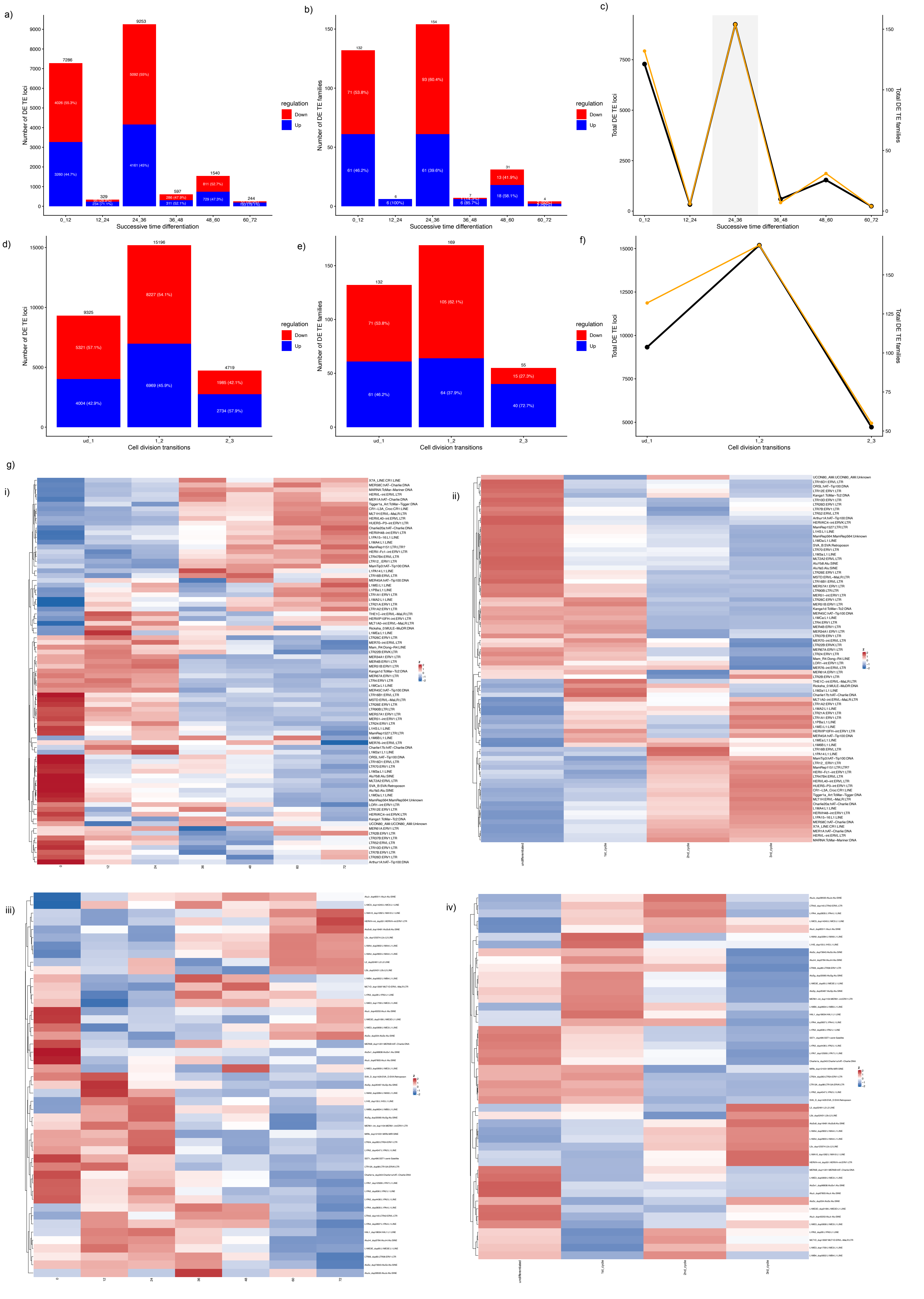
